# Introducing a gastric microbial model community

**DOI:** 10.64898/2026.03.11.710753

**Authors:** Mirjam Dannborg, Sara Lindén, Kaisa Thorell, Johan Bengtsson-Palme

**Author notes:** Correspondence: Johan Bengtsson-Palme Division of Systems and Synthetic biology, Department of Life sciences, Chalmers University of Technology, Kemivägen 10, 412 96, Gothenburg, Sweden.

## Abstract

Persistent colonization by *Helicobacter pylori* in the stomach is connected to the development of different gastric pathologies including gastric cancer. Around half of the world’s population is estimated to be colonized by *H. pylori*, but few develop symptoms. The bacterium dominates the gastric mucosa of individuals who are *H. pylori*-positive by conventional testing, however, low abundances of *H. pylori* have also been detected in individuals who test negative, suggesting low-level colonization The factors driving the predominance of *H. pylori* within the gastric microbiota remain unclear, as are its interactions with the gastric microbiota. Yet these interactions may offer important insights into the organism’s colonization success and its role in disease. In this study, we developed a gastric microbial model community, consisting of five species with high abundance and prevalence in gastric samples based on transcriptional data: *H. pylori*, *Escherichia coli*, *Pseudomonas aeruginosa*, *Streptococcus salivarius,* and *Lactobacillus kalixensis*. We established growth parameters and methods to track species abundance *in vitro*. Our results show that *H. pylori* growth is partly inhibited in pair-wise cultures with all other community members, except with *L. kalixensis.* This inhibition can to a certain extent be explained by resource competition, pH modulation of the medium, and the initial inoculum ratio of the members. The gastric microbial model community will allow dissection of the genetic mechanisms behind *H. pylori* interactions with other members of the gastric microbiota, and how these interactions influence *H. pylori* pathogenesis, providing a non-invasive, low resource model for future studies of the gastric microbiome.

## Introduction

*Helicobacter pylori* is a gram-negative, spiral-shaped bacterium specialized to survive in and colonize the gastric mucosa, an environment long thought to be sterile due to its harsh conditions. *H. pylori* colonization is linked to the development of pathological conditions such as gastritis, peptic ulcer disease, and gastric cancers. Half of the world’s population is estimated to carry *H. pylori*, but only a fraction of individuals develop symptoms (Peleteiro et al., 2014). Why *H. pylori* colonization only causes disease in certain individuals is not yet understood.

*H. pylori* and other bacteria that live in the gastric mucosa together constitute the gastric microbiota. DNA studies of gastric mucosal bacteria generally show a high interindividual variation and relatively low diversity (Aviles-Jimenez et al., 2014; Bik et al., 2006; Delgado et al., 2013). In patients that test positive for *H. pylori* using conventional testing, the gastric microbiota is completely dominated by *H. pylori*. A majority of infected individuals remain asymptomatic but run a risk of developing *H. pylori*-associated gastric diseases. Interestingly, *H. pylori* has also been detected in individuals testing negative for *H. pylori* by conventional tests, but at a much lower abundance (Bik et al., 2006; Maldonado-Contreras et al., 2011; Sorini et al., 2023; Thorell et al., 2017; Wurm et al., 2018), suggesting a low-level colonization. How and under which conditions *H. pylori* would go from seemingly coexisting with the other gastric bacteria, to dominating the gastric mucosa is currently unknown. Understanding *H. pylori’s* interaction with the other gastric bacteria might bring us closer to deciphering the triggers for *H. pylori* pathogenesis.

Previous studies of the gastric microbiota have largely relied on sequencing technologies to identify the species present, which does not provide information about interspecies interactions in the microbial community. A natural microbiota is often very complex and difficult to study *in vitro.* On the other hand, studying a single bacterium is often not representative of how bacteria exist *in vivo.* A way to overcome this limitation is to use microbial model communities (Bengtsson-Palme, 2020). Microbial model communities act as representatives of more complex natural microbial communities, while still being simple enough to study interbacterial interactions *in vitro*. This study aimed to create a bacterial model community representing the gastric microbiota as a tool to study interbacterial interactions between gastric bacteria, with the ultimate goal of furthering the understanding of *H. pylori* colonization in the stomach.

## Materials and Methods

### Media and antibiotic preparation

Ham’s F12 medium pH=7.4 (N6658, Sigma-Aldrich) was adjusted to pH=6.0 with 1 M HCl using a pH probe (Mettler Toledo) and sterile-filtered using a bottle-top 0.22 µm PES filter (Thermo Scientific), when indicated. The bacteria were cultured in Brucella medium (B3051, Sigma-Aldrich), Luria Bertani (LB) medium (L3022, Sigma-Aldrich), Man-Rogosa-Sharp (MRS) medium (69966-500G, Sigma-Aldrich), and Brain-heart infusion (BHI) medium (237500, BD Difco). When indicated, Tween-80 (P4780, Sigma-Aldrich) 1% (v/v), fetal bovine serum 5% (v/v) (FBS; 10270, Gibco), or defibrinated horse blood 5% (v/v) (SR0050C, Thermo Scientific) was added. For making agar plates, 1.5% agar (05040, Sigma-Aldrich) was added. Irgasan (72779-5G-F, Sigma-Aldrich) was diluted in 1 M NaOH, cefsulodin sodium salt hydrate (C8145, Sigma-Aldrich) was diluted in H_2_O, oxolinic acid (O0877-5G, Sigma-Aldrich) was diluted in 0.5 M NaOH, colistin sulfate salt (C4461-100MG, Sigma-Aldrich) was diluted in H_2_O, and trimethoprim (T7883-5G, Sigma-Aldrich) was diluted in DMSO. Vancomycin in DMSO (SBR00001, Sigma-Aldrich) and polymyxin B in H_2_O (92283, Sigma-Aldrich) were purchased as ready-made solutions. Final concentrations of antibiotics are stated in table 1. All antibiotic stocks were stored at -20°C until use, except polymyxin B which was stored at 4°C.

**Table 1.**
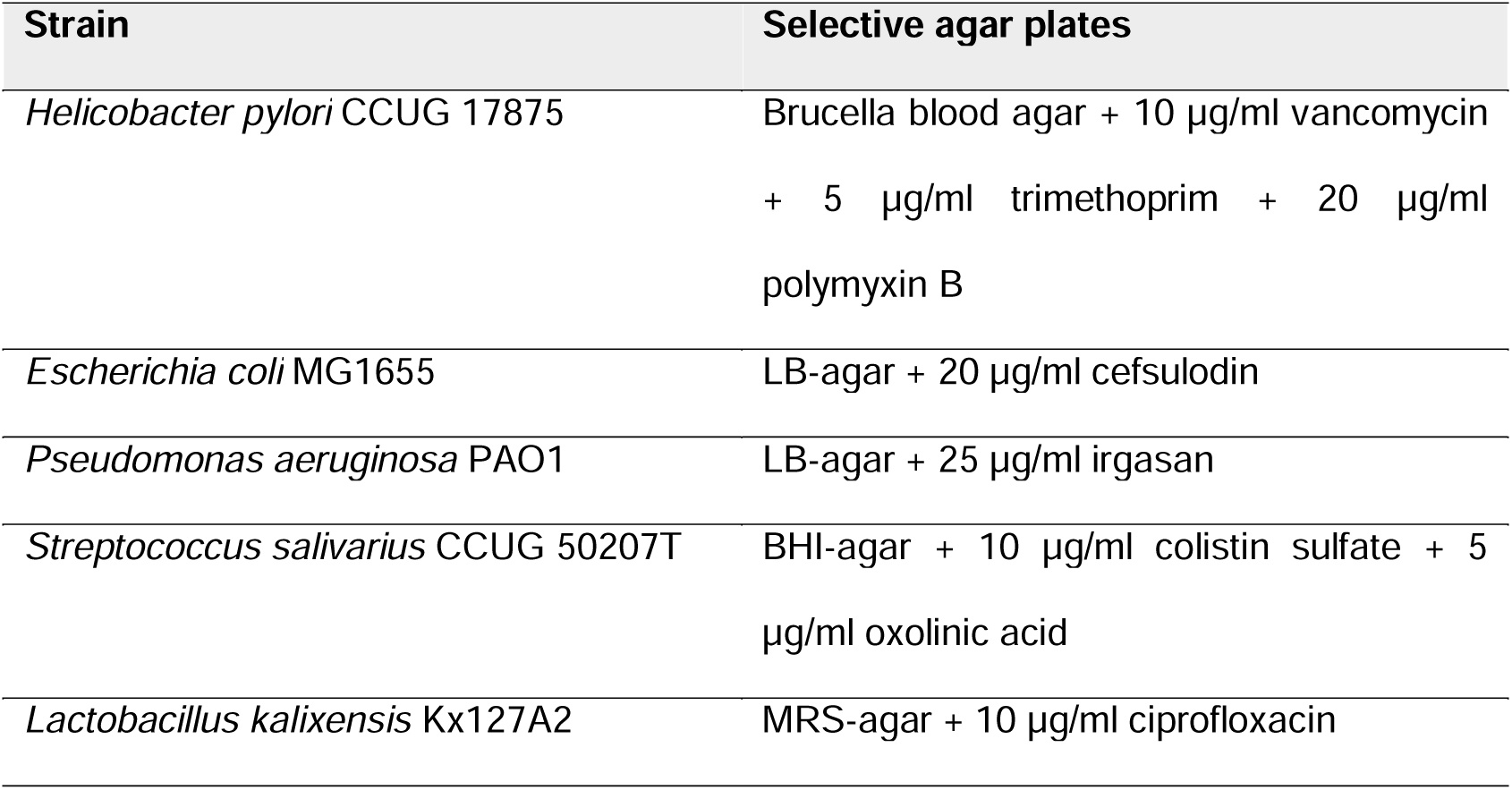
Selective agar for CFU enumeration of the gastric model community.

### Bacterial strains and culture conditions

*Helicobacter pylori* CCUG 17875, *Pseudomonas aeruginosa* PAO1, *Escherichia coli* MG1655, *Streptococcus salivarius* CCUG 50207T, and *Lactobacillus kalixensis* Kx127A2 were acquired from the Culture Collection University of Gothenburg (https://www.ccug.se/). *H. pylori* was cultured on Brucella agar supplemented with 5% (v/v) defibrinated horse blood for 4 days at 37 °C under microaerobic conditions (6-13% O_2_) using a CampyGen pouch (CN0025A, Thermo Scientific). *E. coli* and *P. aeruginosa* were cultured on LB agar at 37 °C over-night. *S. salivarius* was cultured on BHI agar at 37 °C overnight. *L. kalixensis* was cultured on MRS agar with 1% Tween-80 at 37 °C under anaerobic conditions using an AnaeroGen pouch (AN0025A, Thermo Scientific) for 24-48h.

For co-culture experiments, *H. pylori* plates were washed with Brucella broth supplemented with 5% (v/v) FBS and incubated as liquid cultures at 37 °C under microaerobic conditions for 24-48 h, shaking at 150 rpm. For *E. coli* and *P. aeruginosa*, a single colony was resuspended in LB broth and incubated for 16-18 h at 37 °C shaking at 200 rpm. For *S. salivarius*, a single colony was resuspended in BHI broth and incubated at 37 °C for 16-18 h shaking at 200 rpm. For *L. kalixensis*, a single colony was resuspended in MRS broth with 1% (v/v) Tween-80 and stationarily incubated at 37 °C for 24-48 h under anaerobic conditions, in 15 ml polypropylene tubes (525-0604, VWR).

### Species selection

To identify transcriptionally active species in the gastric environment, we reanalyzed data from Thorell et al., 2017 using Metaxa2 (Bengtsson-Palme et al., 2015) with default options. Genus and species classifications were extracted using metaxa2_ttt.

### Co-culture experiments

Before each experiment, the pH of Ham’s F12 medium was adjusted to pH 6.0 using 1 M HCl and sterile-filtered using a bottle-top 0.22 µm PES filter (Thermo Scientific). The pH of the medium was again checked with pH paper (2611-628, Cytiva Whatman) before starting the experiment. Monocultures were adjusted to either OD_600_=0.5 for *H. pylori*, or OD_600_=0.00005 for *E. coli*, *P. aeruginosa*, *S. salivarius*, and *L. kalixensis* by washing overnight cultures in fresh Ham’s F12 pH=6.0. Each strain was diluted 1:5 in a total volume of 200 µl in a flat-bottom 96-well plate (82.1581, Sarstedt), to a final OD_600_ of 0.1 for *H. pylori* and 0.00001 for *E. coli*, *P. aeruginosa*, *S. salivarius*, and *L. kalixensis*. The plates were incubated at 37 °C in microaerobic conditions using a CampyGen pouch for 24h. After each time point, OD_600_ was measured using a VARIOskan LUX plate reader (Thermo Scientific). Subsequently, serial dilutions of the co-cultures were made in sterile 1X PBS, and 5 µl of each dilution was spotted on selective plates (see Table 1) to enumerate colony-forming units (CFUs). *E. coli*, *P. aeruginosa*, and *S. salivarius* selective plates were incubated at 37 °C for 16-18 h. *L. kalixensis* plates were incubated at 37 °C anaerobically for at least 24 h. *H. pylori* selective plates were incubated microaerobically using a CampyGen pouch at 37 °C for 5 days. We found that the *H. pylori*- and *E. coli* selective plates should be made as fresh as possible and preferably stored at 4 °C no longer than 2 weeks before use. The co-cultures were performed in triplicates over three days, and the average CFU/ml was used for plotting.

### Real-time PCR

Primers and probes were designed by aligning the 16S rDNA sequences (retrieved from NCBI, see accession numbers in table 2) of each species using the Clustal Omega Multiple Sequence Alignment tool (Madeira et al., 2024) and Primer-BLAST (Ye et al., 2012). Each probe included three locked nucleic acid bases at unique positions for each species to increase specificity, see table 3 for primer and probe sequences.

**Table 2.** Species and NCBI accession number used for the primer and probe design.

| Species | Accession number |
| --- | --- |
| *Helicobacter pylori* | LS483488 |
| *Escherichia coli* | 4WF1_CA |
| *Pseudomonas aeruginosa* | OQ923798.1 |
| *Streptococcus salivarius* | NR_042776.1 |
| *Lactobacillus kalixensis* | AY253657 |

**Table 3.**
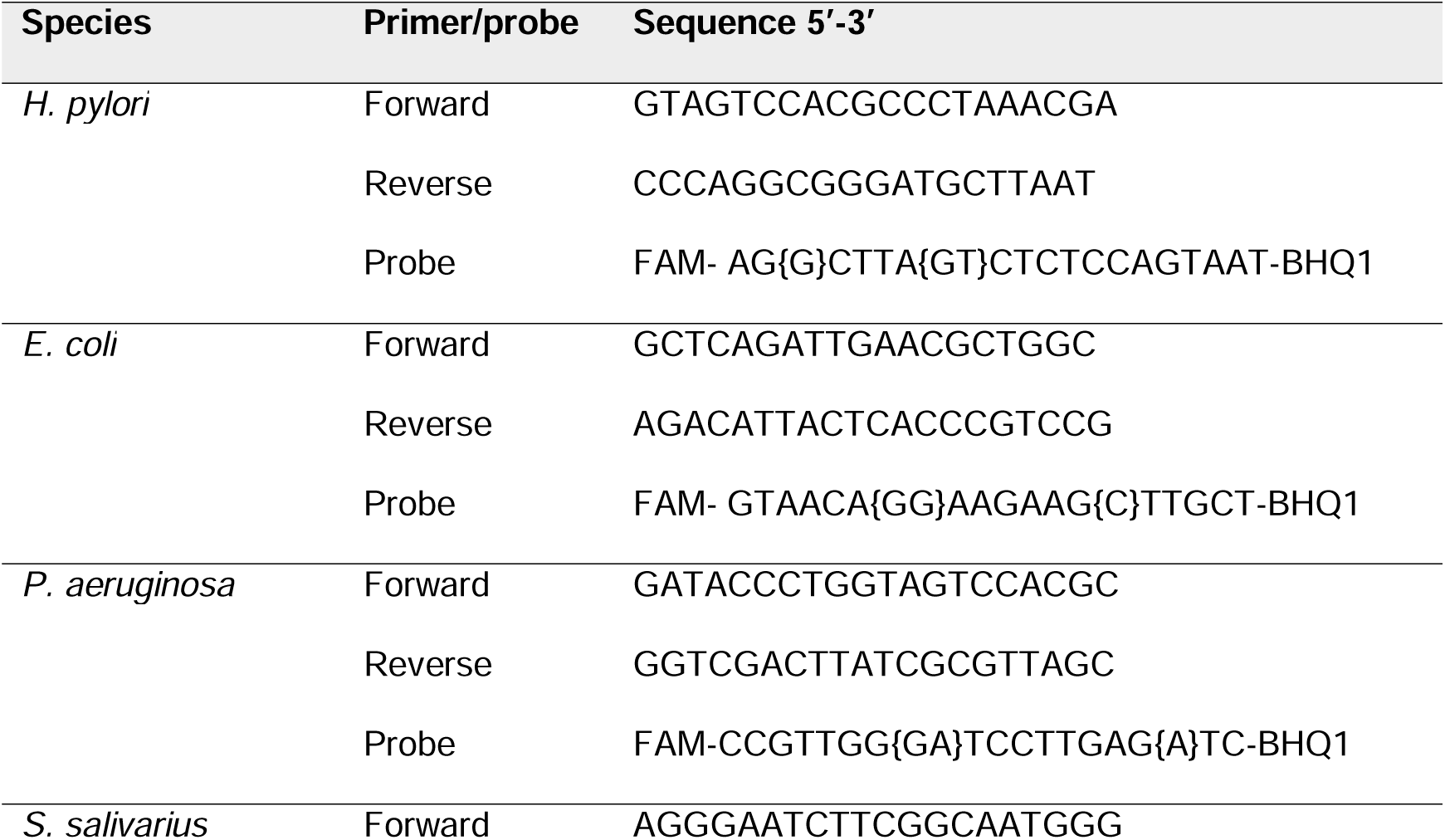
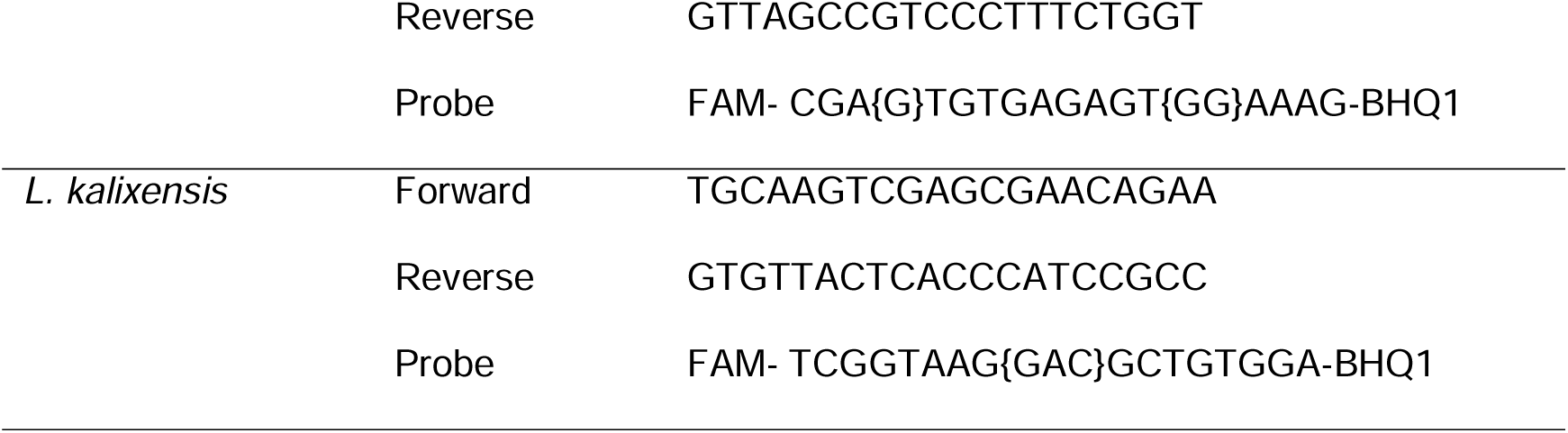
Primers and probe for each of the species in the gastric community. Bases with the {N} brackets are locked nucleic acid bases.

Bacterial samples were first treated with 10 mg/ml lysozyme (Roche) and proteinase K (Qiagen) for 60 min at 37 °C, vortexing for 10 s every 2 min for the first 20 min. DNA was then extracted using the QIAmp DNA mini kit (Qiagen) according to manufacturer’s instructions. The real-time PCR was performed using a Mx-3005 qPCR (Agilent) and the QuantiNova Probe PCR kit (Qiagen) in a 96-well plate using 20 µl reactions according to manufacturer’s instructions. The fast 2-step PCR program was used, as follows: 2 min at 95°C followed by 40 cycles of 6s at 60°C. Three technical replicates were always included, and the cycle threshold was set to 600 after each run.

### Carbon utilization measurement

The carbon source utilization ability of each member was assessed using phenotype microarray plates PM1 and PM2 from BiOLOG according to the manufacturer’s instructions, with the exception that all plates were incubated microaerobically using CampyGen and at 37 °C. Briefly, 100 µl of monoculture cells, diluted in inoculating fluid and redox dye supplied by the manufacturer to a transmittance recommended for each strain, were added to each well of the microarray plate. *E. coli, P. aeruginosa*, *S. salivarius,* and *L. kalixensis* PM plates were incubated for 24 h, and *H. pylori* PM plates were incubated for 48 h, followed by endpoint measurements at 590 nm in a plate reader. The absorbance value for the negative control included in the plates (well A1, containing no carbon source) was subtracted from each well. A cut-off absorbance value was put at 0.1 for us to consider that the bacterium grew using the carbon source. The assays were performed in triplicate over at least two days, and the average value from all replicates of each carbon source was used for calculations.

### Spent medium assay

Monocultures of each strain were adjusted to OD_600_=0.01 in 10 ml of Ham’s F12 (pH 6.0) and incubated at 37 °C under microaerobic conditions with shaking at 140 rpm for 24 h (*E. coli*, *P. aeruginosa*, *S. salivarius*, and *L. kalixensis*) or 48 h (*H. pylori*). The cultures were spun down at 6000 xg for 10 minutes and the supernatants were subsequently sterile filtered using a 0.22 µm PES syringe filter (VWR). The pH of each spent medium was checked again using pH paper (2611-628, Cytiva Whatman) before starting the experiment. For pH-adjusted spent medium, the pH was corrected to pH 6.0 using either HCl or NaOH using a pH-probe and sterile-filtered prior to inoculation. For each species, an overnight culture was washed in sterile PBS and diluted 1/100 in 200 µl of spent medium in a flat-bottom 96-well plate. The plate was incubated for 24h for *E. coli*, *P. aeruginosa*, *S. salivarius*, and *L. kalixensis*, or 48 h for *H. pylori* at 37 °C under microaerobic conditions using a CampyGen pouch.

## Results

### Selecting species for a gastric microbial model community

We sought to construct a gastric microbial model community representative of the mucosal gastric microbiota that can be used as a tool to study interbacterial interactions as well as the molecular mechanisms underlying these interactions. To select relevant species for the model community, we selected species based on consistent transcriptional activity across individuals to increase the likelihood of including bacteria that inhabit the gastric mucosa, as opposed to bacteria that are transiently part of the microbiota. In 2017, Thorell et al. published a study investigating the transcriptional activity of microbial communities in stomach biopsies from individuals with *H. pylori* infection and different levels of gastric pathologies, as well as from healthy controls (Thorell et al., 2017). We reanalyzed the data from this transcriptomics study and among the top genera identified, considering both prevalence and relative abundance, were *Helicobacter*, *Streptococcus*, *Escherichia*, and *Pseudomonas* (Fig. 1A).

**Figure 1.**
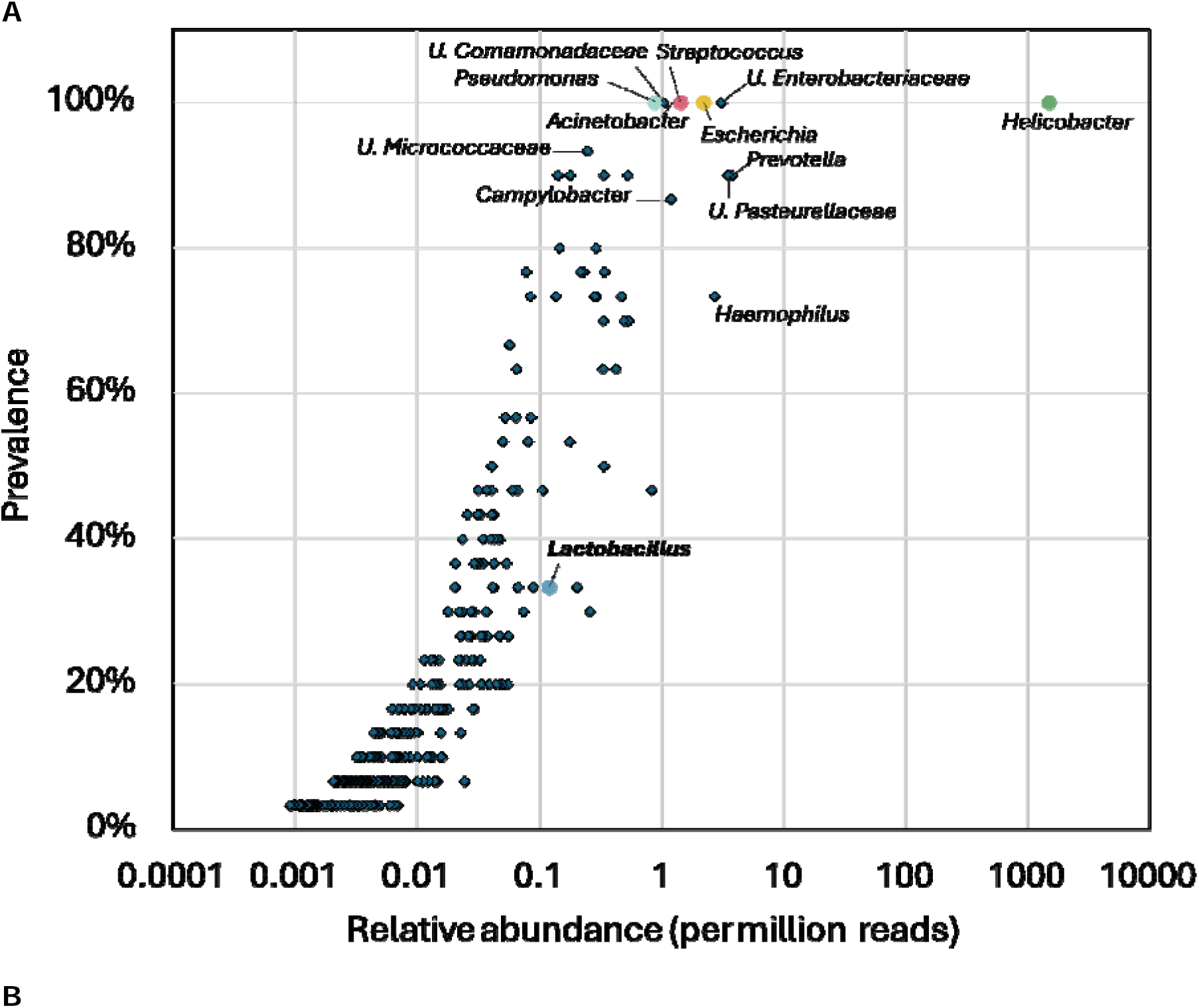
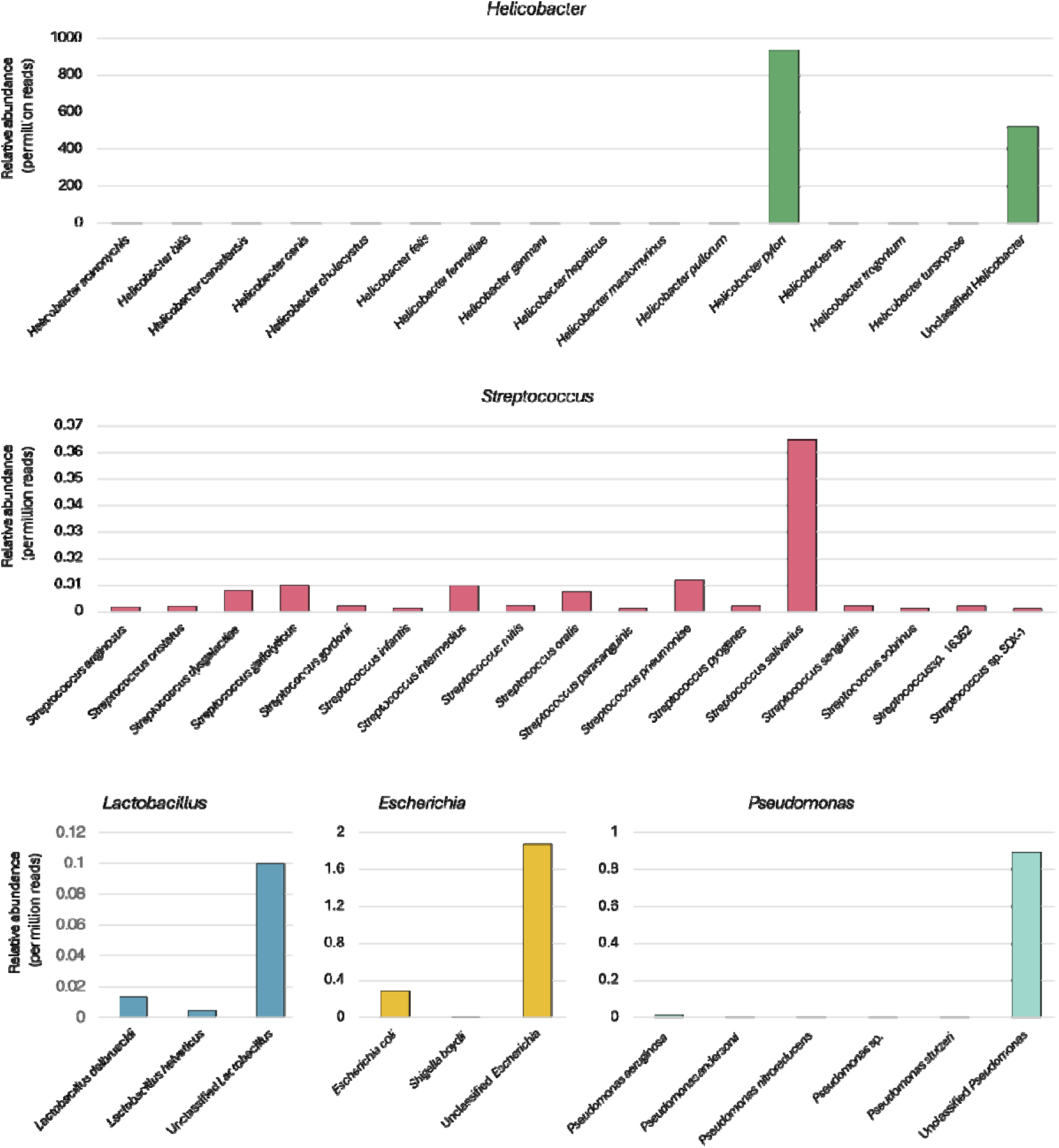
Species and strains selected for the gastric model community. **(A)** Reanalyzed metatranscriptomics data (Thorell et al., 2017) displaying prevalence and relative abundance per million reads across samples for different genera. Colored dots represent the genera chosen to be included in the gastric model community. **(B)** Relative abundance per million reads of species for each of the five genera included in the community.

*Helicobacter* was prevalent in all subjects in the metatranscriptomics study (Fig. 1A), and *H. pylori* was the most abundant species (Fig. 1B). We chose to include *H. pylori* CCUG 17875, an isolate from a gastric antrum biopsy that is well-characterized and has had its genome sequenced. The top species for *Streptococcus* was *Streptococcus salivarius* (Fig. 1B). *S. salivarius* is a core member of the oral- and upper-respiratory tract microbiota (Nakajima et al., 2013) but can also be found in the gastrointestinal tract (Arrieta et al., 2014). It is generally considered a non-pathogenic commensal as it is uncommon in infections. We chose the type-strain *S. salivarius* CCUG 50207T, as it has a full genome sequence available.

Interestingly, few of the rRNA reads in the transcriptomics study identified the genus *Lactobacillus* (Thorell et al., 2017) both in prevalence and abundance (Fig. 1A), even though *Lactobacillus* species have been isolated from both gastric juice and gastric mucosa (Delgado et al., 2013; Roos et al., 2005), as well as identified by culture-independent techniques (Bik et al., 2006; Delgado et al., 2013). For these reasons, we still thought it reasonable to include a *Lactobacillus* species in the model. As we could not identify a species from the transcriptomic data (Fig. 1B), we decided to include *Lactobacillus kalixensis* Kx127A2, an isolate from a healthy gastric mucosa (Roos et al., 2005) with a full genome sequence available. The top species hit for *Escherichia* was unclassified *Escherichia*, followed by *E. coli*. We chose to include *E. coli* MG1655 (K-12 derivate) since the strain is considered non-pathogenic and is well-characterized with a well-annotated genome sequence available.

Most RNA reads assigned to the *Pseudomonas* genus mapped to an unclassified *Pseudomonas* (Fig 1B). We therefore included *P. aeruginosa* in the model community, as it is well-characterized and was found in antral biopsies from individuals with gastritis (Kachuei et al., 2020). The PAO1 strain was selected because it is generally considered less virulent (Grace et al., 2022), and has a fully sequenced genome (Stover et al., 2000).

It is worth noting that the “unclassified” category in the Metaxa2 output includes rRNA sequences that could not be classified due to being too divergent from the reference sequence, and also rRNA sequences that are too similar (or identical) to many species within a genus to be classified to the species level. Thus, we cannot know if, e.g., the unclassified *Pseudomonas* reads in the metatranscriptomes belong to known or novel species of *Pseudomonas*.

Based on the transcriptomic data, we chose to include five species in the gastric model microbial community. The diversity and abundance of the gastric microbiota are relatively low when compared to, for example, the microbiota in the lower intestine, so a five-species model would reflect the typical gastric diversity relatively well. A few other genera were also relatively high in prevalence and abundance, such as *Acinetobacte*r and *Prevotella* (Fig. 1A). However, a five-member community is a manageable size for conducting experiments without using automated solutions, thus we decided to not include more members at this stage. The final gastric microbial model community thus consists of *H. pylori* CCUG 17875, *E. coli* MG1655, *P. aeruginosa* PAO1, *S. salivarius* CCUG 50207T, and *L. kalixensis* Kx127A2 due to their non-pathogenic nature, continuous transcriptional activity in gastric biopsies, and genome sequence availability (Fig. 1C).

### Chemically defined, serum-free, Ham’s F12 supports the growth of all gastric model community members

After finalizing the species and strains to include in the gastric model community, the next step was to find culturing conditions that support growth for all species. The stomach is relatively aerobic with a partial oxygen pressure (pO_2_) of ∼25 mmHg in the gastric lumen of mice (Friedman et al., 2018), but still lower than the atmospheric pO_2_ of ∼160 mmHg. *H. pylori* is a microaerophilic bacterium that requires oxygen for growth but grows poorly under atmospheric oxygen levels. Therefore, the microbial model community was cultured under microaerobic conditions, which was achieved by using CampyGen gas-generating pouches and air-tight containers.

The nutrient requirement for different bacteria varies greatly. Media used for bacterial growth are often complex and nutrient-rich, especially when growing fastidious bacteria such as *H. pylori* that many times also require the addition of serum. Several commonly used rich media were tested, and we found that Elliker broth successfully supported the monoculture growth of all members under microaerobic conditions. However, the number of *P. aeruginosa* colony-forming units (CFUs) after 24h of growth was up to 1 000 000 times larger than for the other members (Suppl. Fig. 1). Earlier studies have shown that Ham’s F12, a chemically defined medium, can support serum-free growth of *H. pylori* (Bessa et al., 2012; Melo et al., 2021; Reynolds & Penn, 1994; Testerman et al., 2001). Using a defined medium allows for the investigation of the metabolic requirements for growth and nutrient competition, among other things. Ham’s F12 is also compatible with some gastric epithelial cell lines. When growing the gastric community members as monocultures in serum-free Ham’s F12 medium, all except *L. kalixensis* had recoverable CFUs after 24h of culture (Fig. 2A). When we cultured all the members together, the fast-growing *P. aeruginosa* and *E. coli* still dominated the community (Fig. 2B) but to a much lesser degree than what was previously seen in a complex medium. Interestingly, the co-culture also revealed community-specific growth support of *L. kalixensis* and *S. salivarius*. (Fig. 2B).

**Figure 2.**
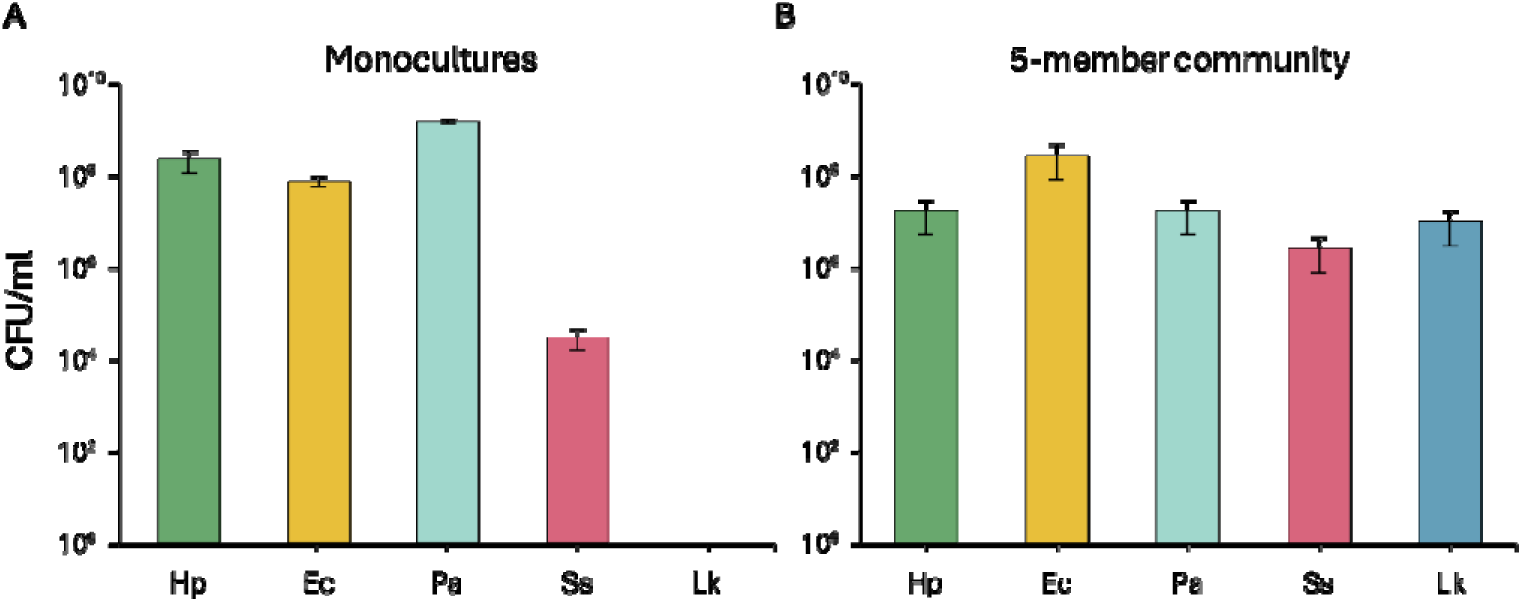
Culturing the gastric model community in Ham’s F12 medium. **(A)** CFUs of the monocultures and **(B)** the five-member community after 24 h of culturing in Ham’s F12 pH 6.0. Hp=*H. pylori*, Ec=*E. coli*, Pa=*P. aeruginosa*, Ss=*S. salivarius*, Lk=*L. kalixensis*. The averages of three independent replicates over three days are plotted. Error bars represent +/-SEM.

When culturing the gastric community together, we initially co-cultured the members in a 1:1 initial inoculum ratio based on OD_600_, but could not recover *H. pylori* CFUs after 24h of co-culture (Suppl. Fig. 2A). Duo-cultures of the members revealed that *E. coli*, *P. aeruginosa*, and *S. salivarius* were the main inhibitors of *H. pylori* growth (Suppl. Fig. 2B). Increasing the initial inoculum ratio to 10 000:1 of *H. pylori* to *E. coli*, *P. aeruginosa,* and *S. salivarius* respectively, rescued *H. pylori* growth without largely affecting the endpoint CFUs of the other members (Suppl. Fig. 2C). Further diluting the other community members did not improve *H. pylori* CFU recovery (Suppl. Fig. 2D). Hence, we settled on an initial inoculum ratio of 10 000:1 of *H. pylori* to the other members.

The gastric milieu has a pronounced pH gradient, ranging from the highly acidic lumen to approaching a neutral pH in proximity to the gastric epithelium. Gastric bacteria, and *H. pylori* in particular, exist in the thick mucus layer protecting the epithelium (Santos et al., 2025), and can therefore be assumed to not be exposed to the luminal pH, but to something closer to a neutral pH. The antral mucosal pH in humans is approximately 5.4 (Quigley & Turnberg, 1987), whereas as a gradient of ∼4.5-7.4 is present in gerbils (Schreiber et al., 2004). With this in mind, we wanted to lower the pH from pH 7.2-7.4 in Ham’s F12 to pH 6.0, to better replicate the *in vivo* setting. All members survived and had recoverable CFUs after 24 h in Ham’s F12 at pH 6.0 (Fig. 2). We also lowered the pH to pH 5.0, but *H. pylori* and *S. salivarius* had no recoverable CFU after 24h (Suppl. Fig. 3).

When co-culturing a microbial community, it is important to be able to track the abundance of the different members over time. Plating dilutions of co-cultures on selective agar plates and subsequent counting of the CFUs will provide insight into the number of viable bacteria remaining in the medium, within the detection limit of the method. We established combinations of selective media that allowed the specific enrichment of each species in the five-member community, see table 1. As a complement, we also designed primers and probes for a qPCR-based detection of DNA from each member in a co-culture, using DNA extracted from a set number of CFUs as the standard curve (Suppl. Fig. 4). The qPCR method overestimates the amount of CFUs for most members (Suppl. Table 1), which we speculate partly can be explained by amplifying DNA from dead bacteria.

In summary, the gastric model community members could all be cultured together in the chemically defined Ham’s F12 medium adjusted to pH 6.0, with an initial inoculum ratio of 10 000:1 of *H. pylori* to each of the other members, with recoverable CFUs after 24 h of co-culture.

The abundance of each member after co-culture can be recorded via CFU enumeration on selective plates or by using a qPCR based method.

### *Helicobacter pylori* only receive negative incoming interactions in duo-cultures

To investigate the interbacterial interactions governing the community structure, we cultured the community in all possible subcommunities, a total of 31 different combinations of mono-, duo-, trio-quartet-, and quintet-cultures (Fig. 2, Suppl. Fig. 5, 6, and 7). Figure 3 shows a network of the duo-culture interaction patterns observed, based on changes in the CFUs compared to monocultures. A negative interaction is defined as a reduced number of CFUs in the duo-culture compared to the monoculture, and a positive interaction as an increase in CFUs in the duo-culture compared to the monoculture. A neutral interaction is defined as no or only small changes in CFUs between the duo-culture and monoculture. The normalized CFU data used to draw the interaction map can be found in Supplemental Table 2.

**Figure 3.**
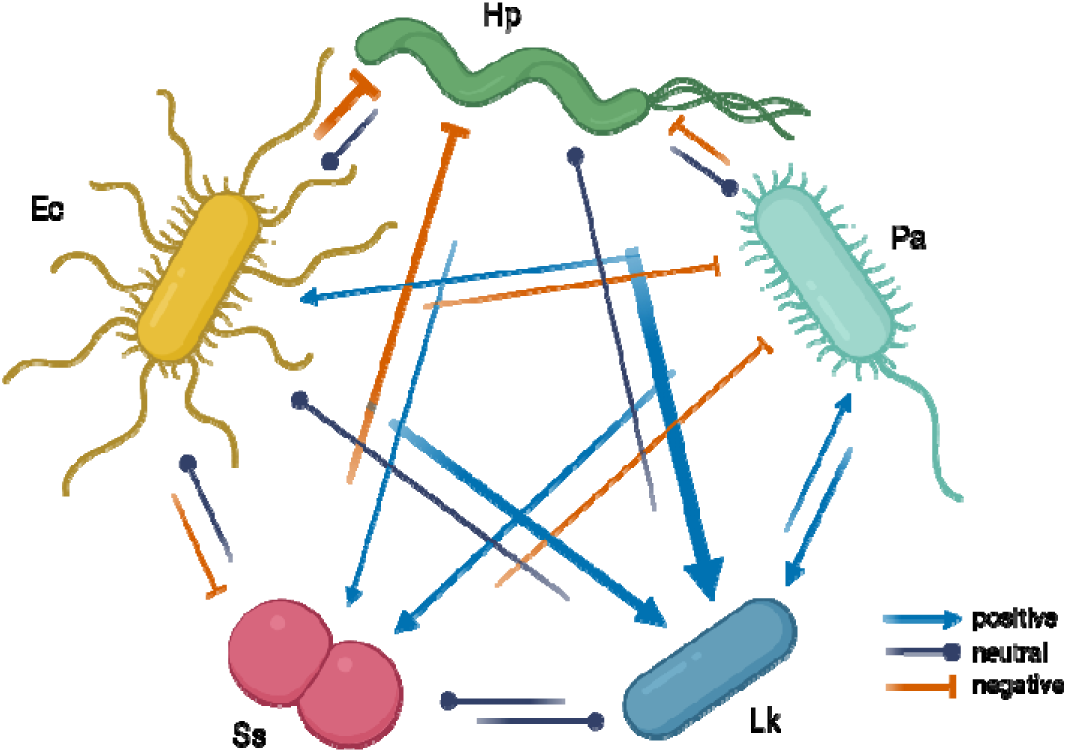
Interaction network based on CFU counts from duo-cultures compared to monocultures. Line thickness is based on the log10 normalized change in CFUs compared to the monoculture CFUs (see Suppl. Table 1) and is indicative of the strength of the interaction. Based on experiments from Supplemental Figure 3 and. Hp=*H. pylori*, Ec=*E. coli*, Pa=*P. aeruginosa*, Ss=*S. salivarius*, Lk=*L. kalixensis*. Created in BioRender. Dannborg, M. (2026) https://BioRender.com/9ldgi2u.

Even with adjusted inoculum ratios, *H. pylori* mainly received negative incoming interactions from the other members, except *L. kalixensis* (Fig. 3). *E. coli* was the greatest inhibitor of *H. pylori,* followed by *S. salivarius* and *P. aeruginosa*. The most striking incoming positive interactions were seen for *L. kalixensis* (Fig. 3), which could not stably grow as a monoculture in Ham’s F12 but flourished in the five-member community (Fig. 2). When examining the duo-cultures, it was evident that *H. pylori, P. aeruginosa*, and *E. coli* supplied the best conditions for *L. kalixensis* growth, however, not to the same extent. *H. pylori* was the major benefactor of *L. kalixensis* growth (Suppl. Fig. 5). Similarly, *S. salivarius* grew better in the presence of *P. aeruginosa* and *H. pylori* (Fig. 3, Suppl. Fig. 5), with the largest growth support seen in the trio-culture (Suppl. Fig. 6).

### Overlap in carbon source utilization partly explains interaction patterns

One of the major interactions between bacteria inhabiting the same space is nutrient competition. To define the nutrient requirements of the five community members, we analyzed the carbon source utilization capability of each strain using phenotype microarray plates. In short, these plates contain a single carbon source per well, and by adding a redox dye together with bacteria, we can investigate whether respiration, and thus growth, take place using this sole carbon source. In total, we screened 190 carbon sources for each strain. In Figure 4, the carbon source overlap between each pair of members is represented, as well as the total number of carbon sources out of the 190 tested that each strain could grow on. Out of the 57 carbon sources *H. pylori* could utilize, 74% and 75% of those were also utilized by *E. coli* and *P. aeruginosa*, respectively (Fig. 4). This indicates that *E. coli* and *P. aeruginosa* to a large extent can occupy the same niche as *H. pylori*, which is also partly reflected in the inhibition seen in duo-cultures (Fig. 3). *S. salivarius* also inhibited *H. pylori* growth in duo-cultures (Fig. 3). However, *S. salivarius* only utilized 23% of the carbon sources *H. pylori* used, indicating that there may be another reason for this inhibition. The *H. pylori* and *S. salivarius* co-culture had a drop in pH from 6 to ∼4-5 after 24 h duo-culture, as measured by pH paper (Suppl. Table 3). Although *H. pylori* can survive in low pH, it prefers a neutral environment, indicating that *S. salivarius* lowering the pH might cause growth stress for *H. pylori*. Similarly, the growth inhibition of *H. pylori* by *E. coli* was strongest out of the pairs, and *E. coli* also lowered the endpoint pH to around 5, suggesting a multifactorial inhibition process via nutrient competition and pH modulation. However, the trio-culture of *H. pylori*, *S. salivarius*, and *L. kalixensis* had an endpoint pH of around 5 (Suppl. Table 3), while the CFU of *H. pylori* was similar to the monoculture (Suppl. Fig. 6), pointing to a more complicated network.

**Figure 4.**
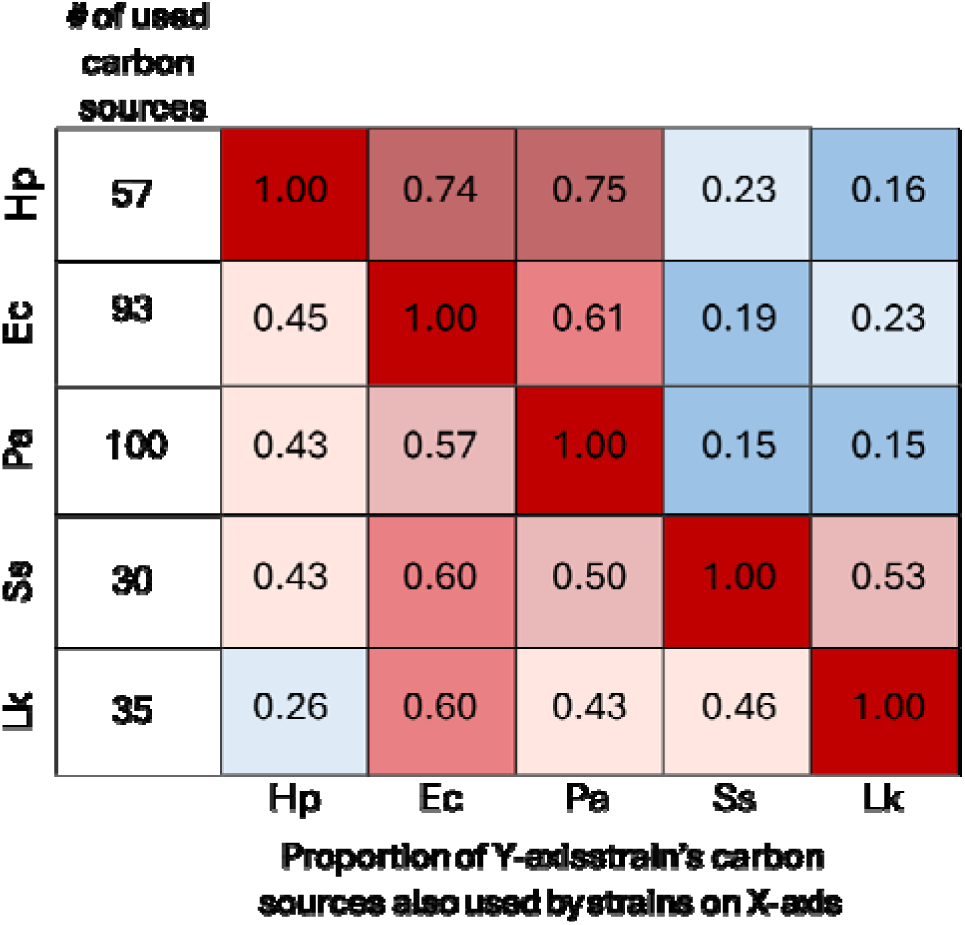
Proportions of carbon source utilization overlap. The total number out of the 190 tested carbon sources a strain can utilize is stated in the leftmost column. The numbers represent the proportion of carbon sources the strains on the X-axis can utilize out of all the carbon sources the strain on the Y-axis utilizes. Hp=*H. pylori*, Ec=*E. coli*, Pa=*P. aeruginosa*, Ss=*S. salivarius*, Lk=*L. kalixensis*. The averages of three independent replicates performed over at least two days were used for calculations.

*H. pylori* and *L. kalixensis* only shared 16% of *H. pylori’s* carbon sources, and 23% of *L. kalixensis’* carbon sources respectively, indicating a low niche overlap between the two, which is in line with the CFU data from the duo-cultures. *L. kalixensis* growth was vastly improved in the presence of *H. pylori,* which could indicate that *H. pylori* provides *L. kalixensis* with previously unattainable nutrients as a by-product of metabolism. The same type of growth support effect was seen for *S. salivarius*. These secondary effects are not captured in these phenotype microarray plate assays, but prompt further metabolic studies of the model community.

Since Ham’s F12 is a defined medium, we could analyze the ability of the members to utilize the carbon sources present in the culturing medium, which were also included in the phenotype microarray plates. 22 out 25 potential carbon sources were included in the phenotype plates. Notably, out of the 10 carbon sources in Ham’s F12 that were tested and that *H. pylori* could utilize, 100% and 90% of those were also utilized by *P. aeruginosa* and *E. coli,* respectively (Suppl. Fig. 8). *L. kalixensis* and *S. salivarius* could only utilize one of the 22 carbon sources tested for in Ham’s F12 (Suppl. Fig. 8), recapitulating the absence of CFUs in *L. kalixensis* monocultures and the relatively poor growth of *S. salivarius* in monoculture. It should be noted that 16 components in Ham’s F12 were not included in the microarray plates (see the full list in Supplemental Table 4). Notably, out of these 16, three components can potentially be used as a carbon source by bacteria: L-cystine, L-tryptophan, and L-tyrosine. The other missing components mostly consist of vitamins, which typically are not used as carbon sources.

Since the observed interactions cannot be explained by nutrient competition or pH modulation alone, we wanted to investigate whether direct cell-to-cell contact was responsible for any growth interaction patterns. We cultured each of the community members in cell-free spent medium from the other members (Fig. 5A, Supple. Table 5)). Overall, similar trends as seen for the co-cultures were observed. *H. pylori* grew poorly in the spent medium of all other members, with *L. kalixensis* spent medium being the least inhibitory. *L. kalixensis* grew better in spent medium from *H. pylori* followed by spent medium from *E. coli.* Interestingly, the positive growth effect from *P. aeruginosa* was not seen when *L. kalixensis* was grown in the spent medium from *P. aeruginosa*. The largest positive effect was seen on *S. salivarius* cultivated in *H. pylori* spent medium, which resulted in a 26 times increase in *S. salivarius* CFUs, compared to the duo-culture of *S. salivarius* and *H. pylori* that resulted in 11 times more *S. salivarius* CFUs (Suppl. Table 1). However, this effect can be, at least partially, due to changes of the medium pH. Both *E. coli* and *S. salivarius* lower the pH of the medium, which will affect the other members based on the pH effects alone.

**Figure 5.**
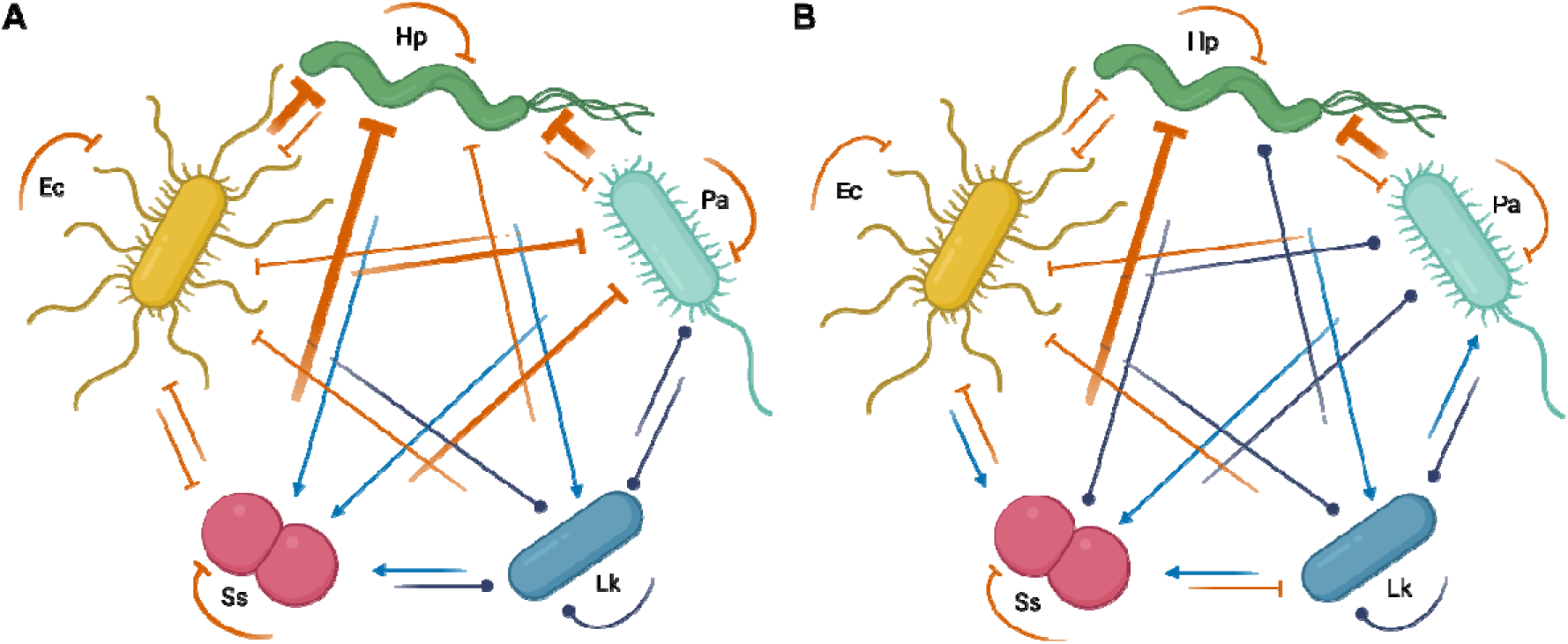
Interaction map representing change in CFU after growth in cell-free spent medium. The change in growth of each community member after growth in (A) cell-free spent medium of the other members or (B) in cell-free spent medium adjusted to pH=6, compared to growth in fresh Ham’s F12 pH 6.0. Line thickness is based on the log10 normalized change in CFUs compared to growth in fresh F12 medium (Suppl. Table 3 and Suppl. Table 4). Hp=*H. pylori*, Ec=*E. coli*, Pa=*P. aeruginosa*, Ss=*S. salivarius*, Lk=*L. kalixensis*. The averages of three independent replicates performed over three days were used for calculations. Created in BioRender. Dannborg, M. (2026) https://BioRender.com/mv9cia8.

To account for potential pH-dependent inhibitory effects, the pH of the spent medium was adjusted to pH 6.0 prior to inoculation (Fig. 5B, Suppl. Table. 6). Following pH-adjustment, several interactions previously classified as negative became neutral. As an example, the inhibitory effects of the *E. coli* spent medium was markedly reduced for most community members after adjusting pH. In contrast, the *S. salivarius* and *P. aeruginosa* spent medium still had a strong inhibitory effect on *H. pylori* growth, despite the higher pH. These findings suggest that inhibitory mechanisms beyond acid stress may contribute to the observed inhibitory effect, although their nature and relative importance remain to be elucidated.

## Discussion

In this paper, we present a five-species microbial model community representing the gastric microbiota that can be used to study interactions between *H. pylori* and other stomach-associated bacteria. We showed that all five species can grow together in the chemically defined, serum-free Ham’s F12 medium. When cultured together, the bacteria influenced each other’s growth and *H. pylori* received predominantly negative incoming interactions in duocultures. The observed interaction dynamics can be at least partly attributed to overlapping carbon-source utilization.

Resource competition is indeed a key mechanism underlying negative interactions in microbial communities and can be used as a functional measure of niche overlap. In our system, *E. coli* and *P. aeruginosa* utilized a majority of carbon sources used by *H. pylori* and is likely the basis of the negative growth interactions between the bacteria. Niche overlap is a major contributing factor to the level of colonization resistance an established microbiota can excerpt to invading pathogens or overgrowth of indigenous opportunists (Sorbara & Pamer, 2019). This was recently demonstrated in microbial communities by Spragge et al., 2023, who showed that key species that consume the same nutrients as the invader offered protection against the establishment and survival of pathogens within the community. Another study similarly showed that a rationally designed low-complexity microbial community was efficient in providing colonization resistance against the gut pathogen *Clostridium difficile,* partly due to resource competition (Hromada et al., 2021). Colonization resistance of a community can also depend on prior disturbances. Pre-treatment with non-antibiotic drugs has been shown to disrupt a bacterial model community and facilitate pathogen invasion (Grießhammer et al., 2023). This is also reflected *in vivo*, where regular and prolonged intake of, e.g., proton-pump inhibitors (Cheung et al., 2018) and nonsteroidal anti-inflammatory drugs (NSAIDs) (Sostres et al., 2014) have been connected to *H. pylori* symptomatic infection in patients.

In our model community, *H. pylori* was negatively impacted by growing with most members in the gastric community, except *L. kalixensis.* This was unexpected, since *Lactobacillus* species tend to be viewed as one of the most promising probiotic bacteria against *H. pylori* colonization. Probiotics strains of *Lactobacillus* in combination with classical *H. pylori* eradication therapy have shown to reduce the side effects of therapy (such as diarrhea) and overall improve eradication results, in randomized controlled trials (Yang et al., 2024). *L. kalixensis* could still prevent, for example, the adhesion of *H. pylori* to gastric epithelial cells, which we have not tested for here. Nevertheless, this highlights that probiotic abilities, and in extension species interactions, are species- and even strain-dependent (Yang et al., 2024). In our gastric microbial model community, the selected strains are all considered to be either non-pathogenic or low virulence. However, most of the included species encompass other strains with pathogenic or otherwise different phenotypes *in vivo*, allowing relatively easy strain exchanges in the gastric community in future studies of specific features, including, for example, more virulent *H. pylori* strains to investigate potential differences in colonization and competition with other members.

Microbial model communities allow mechanistic studies of community-specific traits that are overlooked in monospecies studies, and other microbial model communities of varying complexity have been constructed to represent different human-associated microbiota. A 12-member microbial model community was assembled to represent the main phyla of the human colonic microbiota and demonstrated that duocultures was the main driver to determine higher-order interactions in microbial communities (Venturelli et al., 2018). A nine-species microbial model community representing the human skin microbiome could successfully coculture anoxic and oxic bacterial species commonly found on human skin, and replicate results from human cosmetic trials *in vitro* using the model community (Lekbua et al., 2024). A four-member microbial model community representing the cystic fibrosis lung microbiota showed community-specific increase in antibiotic tolerance of *P. aeruginosa* (Jean-Pierre et al., 2023). The same microbial model community was also used to identifying cross-feeding mechanisms allowing survival of members not able to survive in monoculture (El Hafi et al., 2024). Microbial model communities will in the future also allow more complex studies of microbiome-host interactions using e.g. organ-on-a-chip models and 3D cell-cultures such as organoids. Our microbial model community enables similar studies of the gastric ecosystem and provide a foundation for identifying important interspecies interactions between the stomach-associated bacteria.

Indeed, *H. pylori* colonizing the gastric mucosa is a multi-factorial process and there is an interplay between the host, the environment, and the gastric microbiota that is not captured by studying a microbial model community alone. As mentioned above, non-antibiotic drugs such as proton pump inhibitors and NSAIDs induce environmental changes in the stomach that may favor *H. pylori* growth. The genomic background of *H. pylori* strains also matters, with a pronounced population structure and genetic variability. *H. pylori* can carry the Cag (cytotoxin-associated gene) pathogenicity island, which is linked to the development of gastric disease (Noto & Peek, 2012). Other notable virulence factors include the BabA and SabA adhesins, which facilitate *H. pylori* binding to mucins (Aspholm-Hurtig et al., 2004; Lindén et al., 2002) and epithelial surfaces (Borén et al., 1993), and the cytotoxin VacA (Leunk et al., 1988), which all have been implicated in worsened gastric pathologies (Doohan et al., 2021; Palframan et al., 2012). In addition, *H. pylori* has many strategies to evade both the innate- and adaptive immune system (Fan et al., 2024). Host gene polymorphisms can further facilitate *H. pylori* colonization, such as in changes to cytokines like IL1-β and TNF-α, which regulate acid secretion in the stomach, and polymorphisms in immune receptors such as toll-like receptors and NOD-like receptors (Clyne & Rowland, 2019).

Although the gastric microbial model community and the culturing conditions presented here do not precisely recapitulate the host environment these bacteria exist in *in vivo*, the use of model community will allow us to discern larger and transferable patterns. Furthermore, the ability to exchange strains in the community and follow how this changes community properties allow contextualizing the host-pathogen interactions and what mediates successful colonization of the human stomach. Such information can be used to refine hypotheses before testing in more complex systems, such as animal models (Bengtsson-Palme, 2020).

In summary, we here present a first gastric bacterial model community together with culture conditions and profiling of carbon utilization of the community members as a resource for the research community to investigate *H. pylori* interactions with other gastric bacteria. We anticipate that this system will serve as a foundation for further studies of host-microbe interactions in the gastric environment.

## Acknowledgments

We thank Emil Burman for helpful discussion during the study and Tora Hulterström for helpful discussions and feedback of the manuscript.

This work was supported by the Swedish Research Council (VR) under grant 2020-03629; the Data-Driven Life Science (DDLS) program supported by the Knut and Alice Wallenberg Foundation under grant KAW 2020.0239; and the Swedish Foundation for Strategic Research under grant FFL21-0174.

## Data availability

The RNA-seq data is available in the ArrayExpress public repository (www.ebi.ac.uk/arrayexpress) under accession number E-MTAB-3689.

