## Supplemental material for "Introducing a gastric microbial model community"

**Supplemental table 1.** Actual CFU from culturing and calculated CFU from qPCR Ct-values from full community co-culture. The CFU/ml are averages of independent three experiments performed over three days.

| **Species** | **CFU/ml from culturing** | **Estimated CFU/ml from qPCR** |
| --- | --- | --- |
| *H. pylori* | 1.54E+06 | 1.94E+07 |
| *E. coli* | 2.08E+08 | 8.94E+07 |
| *P. aeruginosa* | 2.02E+08 | 3.45E+09 |
| *S. salivarius* | 9.93E+06 | 6.82E+08 |
| *L. kalixensis* | 5.54E+06 | 4.40E+05 |

**Supplemental table 2.** Log-transformed CFU data from duo-cultures normalized to respective monoculture used to visualize the interaction network in Figure 3. N=neutral (no change in CFU), (+)=at least 0.4 log10 increase in CFU, (-)=at least 0.4 log10 decrease in CFU. Based on experiments performed in triplicates over three different days. Hp=*H. pylori*, Ec=*E. coli*, Pa=*P. aeruginosa*, Ss=*S. salivarius*, Lk=*L. kalixensis.*

| **Culture** | **Strain** | **#CFU in duoculture/**  **#CFU in monoculture** | **Log10 of CFU ratio** | **Interpretation** |
| --- | --- | --- | --- | --- |
| Hp+Ec | Hp | 0.0002 | -3.913 | ---- |
|  | Ec | 1.0914 | -0.024 | N |
| Hp+Pa | Hp | 0.1270 | -1.158 | - |
|  | Pa | 0.8019 | -0.018 | N |
| Hp+Ss | Hp | 0.0020 | -3.003 | --- |
|  | Ss | 11.5601 | 1.0275 | + |
| Hp+Lk | Hp | 2.0322 | 0.0287 | N |
|  | Lk | 8125000.0000 | 6.7936 | +++++++ |
| Ec+Pa | Ec | 3.6548 | 0.5266 | + |
|  | Pa | 0.0527 | -0.736 | - |
| Ec+Ss | Ec | 0.9340 | -0.062 | N |
|  | Ss | 0.1176 | -1.091 | - |
| Ec+Lk | Ec | 0.8731 | -0.017 | N |
|  | Lk | 7500.5000 | 3.7118 | ++++ |
| Pa+Ss | Pa | 0.1118 | -0.493 | - |
|  | Ss | 115.6010 | 2.0444 | ++ |
| Pa+Lk | Pa | 2.9827 | 0.4316 | + |
|  | Lk | 225.0000 | 2.301 | ++ |
| Ss+Lk | Ss | 0.1269 | -0.154 | N |
|  | Lk | 1.0000 | 0 | N |

**Supplemental table 3.** **End-point pH of the different co-cultures.** The initial pH of Ham’s F12 was adjusted to pH=6.0 as measured by a pH probe (Mettler Toledo). The endpoint pH was checked using pH paper (Whatman Cytiva) after 24h of culture. H=*H. pylori*, E=*E. coli*, P=*P. aeruginosa*, S=*S. salivarius*, L=*L. kalixensis*. Blank=medium only control.

| **Culture** | **Endpoint pH** |
| --- | --- |
| H | 7.0 |
| E | 4.0-5.0 |
| P | 6.0-7.0 |
| S | 5.0-6.0 |
| L | 6.5-7.0 |
| HE | 5.0-6.5 |
| HP | 6.0-6.5 |
| HS | 5.0-6.0 |
| HL | 7.0 |
| EP | 6.0-6.5 |
| ES | 4.5-5.0 |
| EL | 4.5-5.0 |
| PS | 6.0-6.5 |
| PL | 6.0-6.5 |
| SL | 5.0-6.0 |
| HEP | 6.5-7.0 |
| HES | 5.0 |
| HEL | 5.0-6.5 |
| HPS | 6.5 |
| HPL | 6.5-7.0 |
| HSL | 5.0-5.5 |
| EPS | 6.5-7.0 |
| EPL | 6.5-7.0 |
| ESL | 5.0-6.0 |
| PSL | 6.0-7.0 |
| HEPS | 6.0-6.5 |
| HEPL | 6.0-6.5 |
| HESL | 4.5-5.5 |
| HPSL | 6.0-6.5 |
| EPSL | 6.0-6.5 |
| HEPSL | 6.0-6.5 |
| Blank | 6.0-6.5 |

**Supplemental table 4.** Components of Ham’s F12 not included in the phenotype microarray plates PM1 and PM2 (BiOLOG).

| **Component** | **Role** |
| --- | --- |
| L-cystein | Amino acid |
| L-tryptophan | Amino acid |
| L-tyrosine | Amino acid |
| D-Biotin | Vitamin (B_7_) |
| Choline Chloride | Ammonium salt |
| Folic acid | Vitamin (B_9_) |
| Niaciamid | Vitamin (B_3_) |
| D-Panthothenic acid | Vitamin (B_5_) |
| Pyridoxine | Vitamin (B_6_) |
| Riboflavin | Vitamin (B_2_) |
| Thiamine | Vitamin (B_1_) |
| Vitamin B_12_ | Vitamin (B_12_) |
| Hypoxanthine | Purine derivative, nitrogen source |
| Linoleic acid | Omega-6 fatty acid |
| Phenol red | pH indicator |
| Thioctic acid | Organosulfur compound, enzymatic co-factor |

**Supplemental table 5.** Log-transformed CFU data from monocultures grown for 24 h in indicated spent medium normalized to monoculture grown for 24h in fresh medium, used to visualize the interaction network in Figure 5A. N=neutral (no change in CFU), (+)=at least 0.4 log10 increase in CFU, (-)=at least 0.4 log10 decrease in CFU. Based on experiments performed in triplicates over three different days. Hp=*H. pylori*, Ec=*E. coli*, Pa=*P. aeruginosa*, Ss=*S. salivarius*, Lk=*L. kalixensis.*

| **Culture** | **Medium** | **#CFU in spent F12/** | **Log10 of CFU ratio** | **Interpretation** |
| --- | --- | --- | --- | --- |
|  |  | **#CFU in fresh F12** |  |  |
| Hp | Fresh F12 | 1 | 0 | N |
|  | Hp spent | 0.014920513 | -1.826216247 | -- |
|  | Ec spent | 3.00135E-07 | -6.522682739 | ------- |
|  | Pa spent | 3.00135E-07 | -6.522682739 | ------- |
|  | Ss spent | 3.00135E-07 | -6.522682739 | ------- |
|  | Lk spent | 0.299818211 | -0.523141992 | - |
| Ec | Fresh F12 | 1 | 0 | N |
|  | Hp spent | 0.062432432 | -1.204589744 | - |
|  | Ec spent | 0.017027027 | -1.768861175 | -- |
|  | Pa spent | 0.070945946 | -1.149072416 | - |
|  | Ss spent | 0.042297297 | -1.373687382 | - |
|  | Lk spent | 0.208108108 | -0.681710999 | - |
| Pa | Fresh F12 | 1 | 0 | N |
|  | Hp spent | 0.033382625 | -1.476479519 | - |
|  | Ec spent | 0.00057671 | -3.239042671 | --- |
|  | Pa spent | 0.025397412 | -1.595210532 | -- |
|  | Ss spent | 0.000438688 | -3.357844626 | --- |
|  | Lk spent | 0.426987061 | -0.369585285 | N |
| Ss | Fresh F12 | 1 | 0 | N |
|  | Hp spent | 26.10182371 | 1.416670852 | + |
|  | Ec spent | 0.094984802 | -1.022345876 | - |
|  | Pa spent | 3.433510638 | 0.535738397 | + |
|  | Ss spent | 0.010258359 | -1.988922121 | -- |
|  | Lk spent | 5.801291793 | 0.76352471 | + |
| Lk | Fresh F12 | 1 | 0 | N |
|  | Hp spent | 6.136022514 | 0.787886944 | + |
|  | Ec spent | 1.200750469 | 0.079452765 | N |
|  | Pa spent | 0.876547842 | -0.057224375 | N |
|  | Ss spent | 0.770637899 | -0.113149637 | N |
|  | Lk spent | 0.407129456 | -0.390267475 | N |

**Supplemental table 6.** Log-transformed CFU data from monocultures grown for 24 h in indicated pH-adjusted spent medium normalized to monoculture grown for 24h in fresh medium, used to visualize the interaction network in Figure 5B. N=neutral (no change in CFU), (+)=at least 0.4 log10 increase in CFU, (-)=at least 0.4 log10 decrease in CFU. Based on experiments performed in triplicates over three different days. Hp=*H. pylori*, Ec=*E. coli*, Pa=*P. aeruginosa*, Ss=*S. salivarius*, Lk=*L. kalixensis*

| **Culture** | **Medium** | **#CFU in spent, pH-adjusted F12/** | **Log10 of CFU ratio** | **Interpretation** |
| --- | --- | --- | --- | --- |
|  |  | **#CFU in fresh F12** |  |  |
| Hp | Fresh F12 | 1 | 0 | N |
|  | Hp spent | 0.032491736 | -1.48822708 | - |
|  | Ec spent | 0.216059685 | -0.66542626 | - |
|  | Pa spent | 4.17537E-07 | -6.37930552 | ------ |
|  | Ss spent | 4.17537E-07 | -6.37930552 | ------ |
|  | Lk spent | 1.69968138 | 0.230367517 | N |
| Ec | Fresh F12 | 1 | 0 | N |
|  | Hp spent | 0.038796296 | -1.41120973 | - |
|  | Ec spent | 0.034259259 | -1.46522203 | - |
|  | Pa spent | 0.117592593 | -0.92962003 | - |
|  | Ss spent | 0.153703704 | -0.81331567 | - |
|  | Lk spent | 0.275 | -0.56066731 | - |
| Pa | Fresh F12 | 1 | 0 | N |
|  | Hp spent | 0.162269805 | -0.78976229 | - |
|  | Ec spent | 0.93932862 | -0.02718244 | N |
|  | Pa spent | 0.119641495 | -0.92211817 | - |
|  | Ss spent | 2.292212642 | 0.360254903 | N |
|  | Lk spent | 2.558969416 | 0.408065095 | + |
| Ss | Fresh F12 | 1 | 0 | N |
|  | Hp spent | 0.941997063 | -0.02595045 | N |
|  | Ec spent | 11.49045521 | 1.060337234 | + |
|  | Pa spent | 5.528634361 | 0.742617869 | + |
|  | Ss spent | 0.243039648 | -0.61432287 | - |
|  | Lk spent | 12.29809104 | 1.089837704 | + |
| Lk | Fresh F12 | 1 | 0 | N |
|  | Hp spent | 3.835714286 | 0.58384625 | + |
|  | Ec spent | 0.81282266 | -0.0900042 | N |
|  | Pa spent | 0.710344828 | -0.14853078 | N |
|  | Ss spent | 0.320197044 | -0.49458268 | - |
|  | Lk spent | 0.5591133 | -0.25250018 | N |

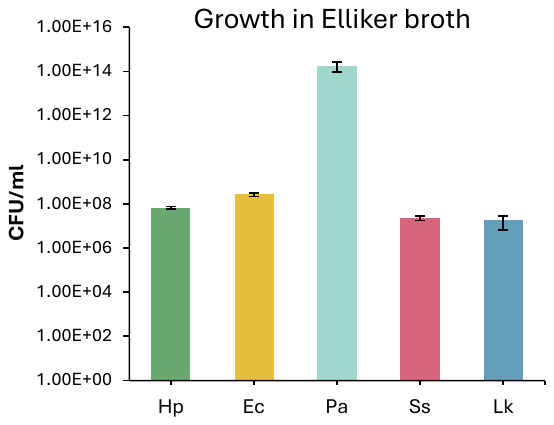

**Supplemental Figure.** **1.** CFUs of gastric model community members in Elliker broth. Hp=*H. pylori*, Ec=*E. coli*, Pa=*P. aeruginosa*, Lk=L. *kalixensis*. Performed in triplicates over three days. Error bars represent +/- SEM.

**A**

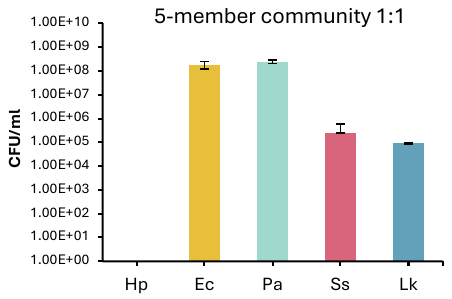

**B**

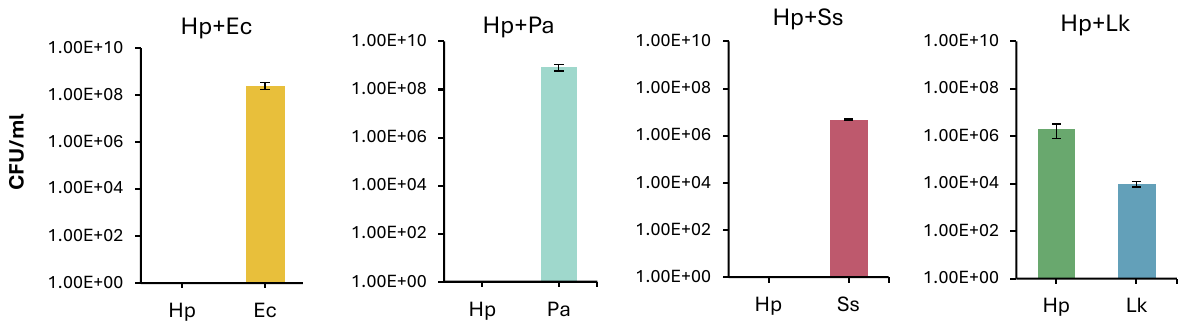

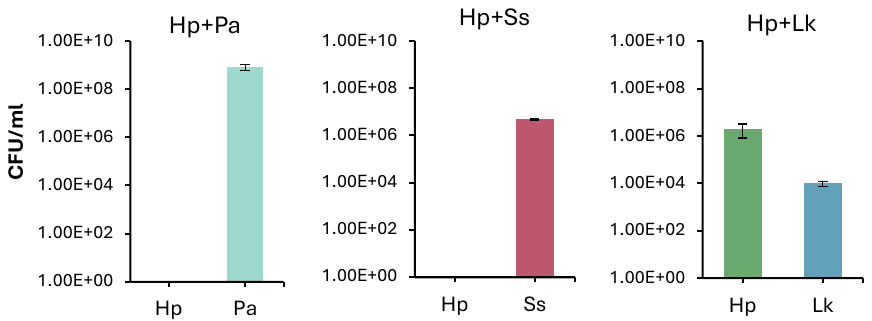

**C**

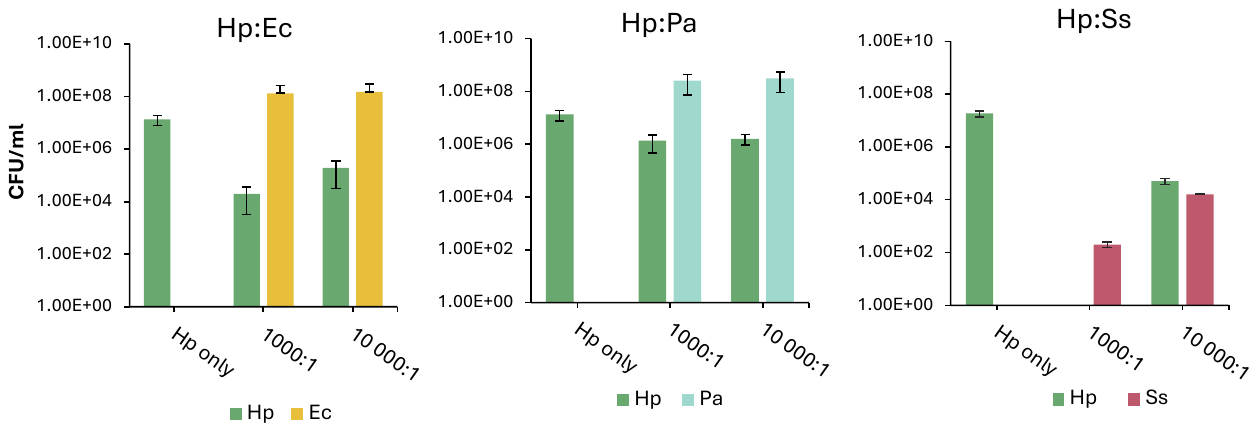

**D**

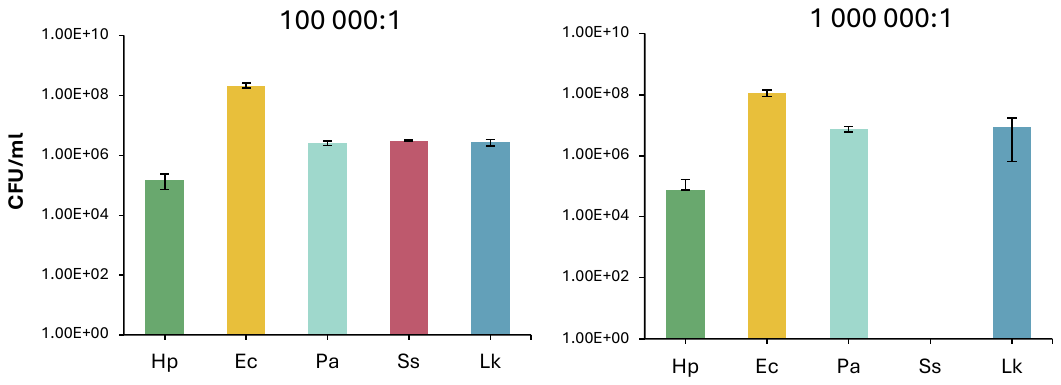

**Supplemental Figure 2. Initial inoculum ratio determines *H. pylori* survival in a community setting (A)** CFUs of the five-member community after 24h co-culture with 1:1 initial inoculum ratio based on OD_600_ **(B)** CFUs of duo-cultures of *H. pylori* with the other community members using a initial inoculum ratio of 1:1 based on OD_600_ **(C)** Duo-cultures of *H. pylori* with *E. coli*, *P. aeruginosa* and *S. salivarius* with an initial inoculum ratio of 1000:1 and 10 000:1 of *H. pylori* to the other members based on OD_600_. **(D)** CFUs of five-member community with an initial inoculum ratio of 100 000:1 and 1 000 000:1 of *H. pylori* to the other members. All co-cultures were done in Ham’s F12 for 24h. Plotted are the averages of experiments performed in duplicate or triplicate on separate days. Error bars represent +/- SD.

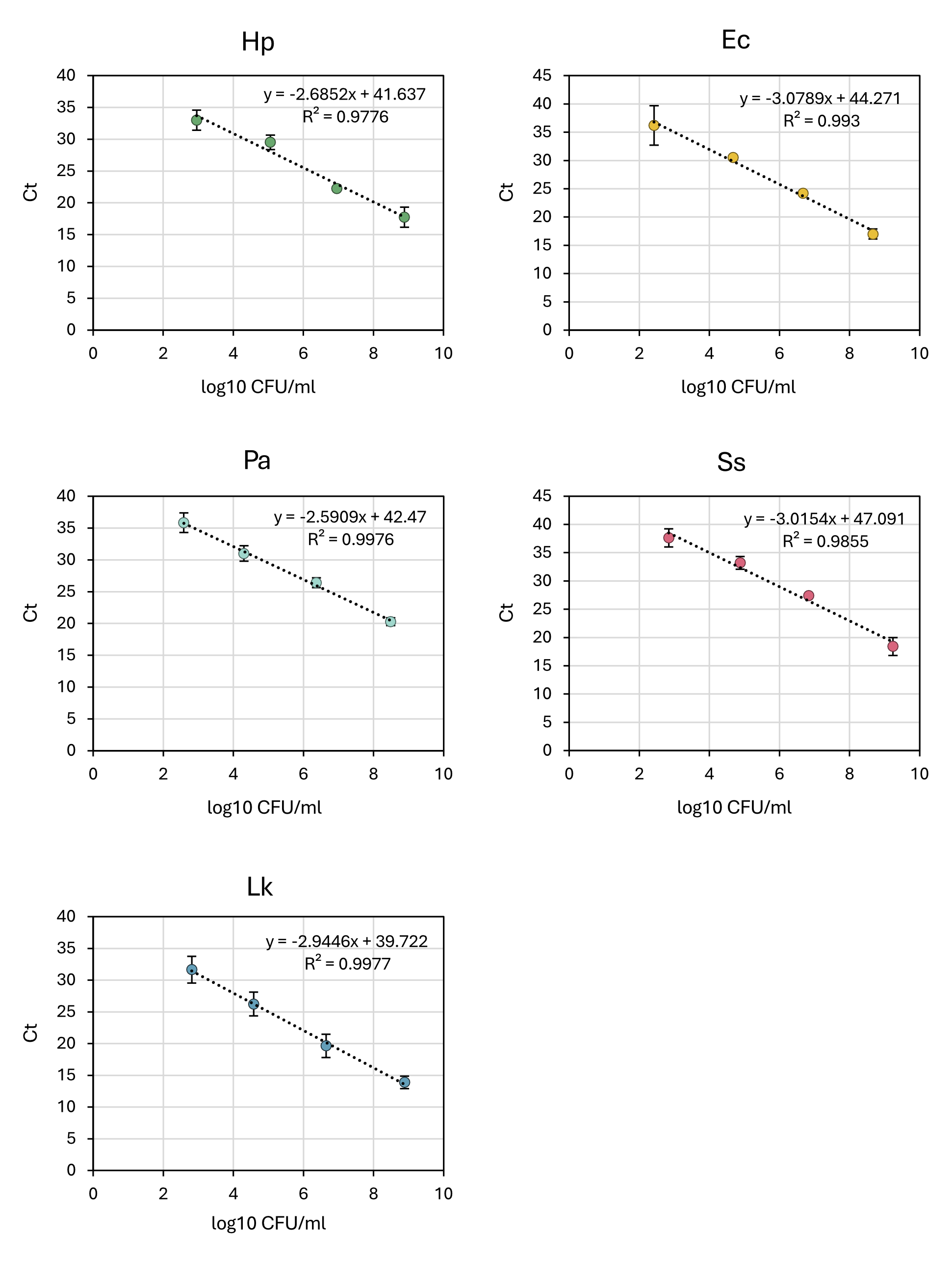

**Supplemental Figure 3. qPCR standard curves relating log10 CFU/ml to Ct for each of the members in the gastric microbial community.**

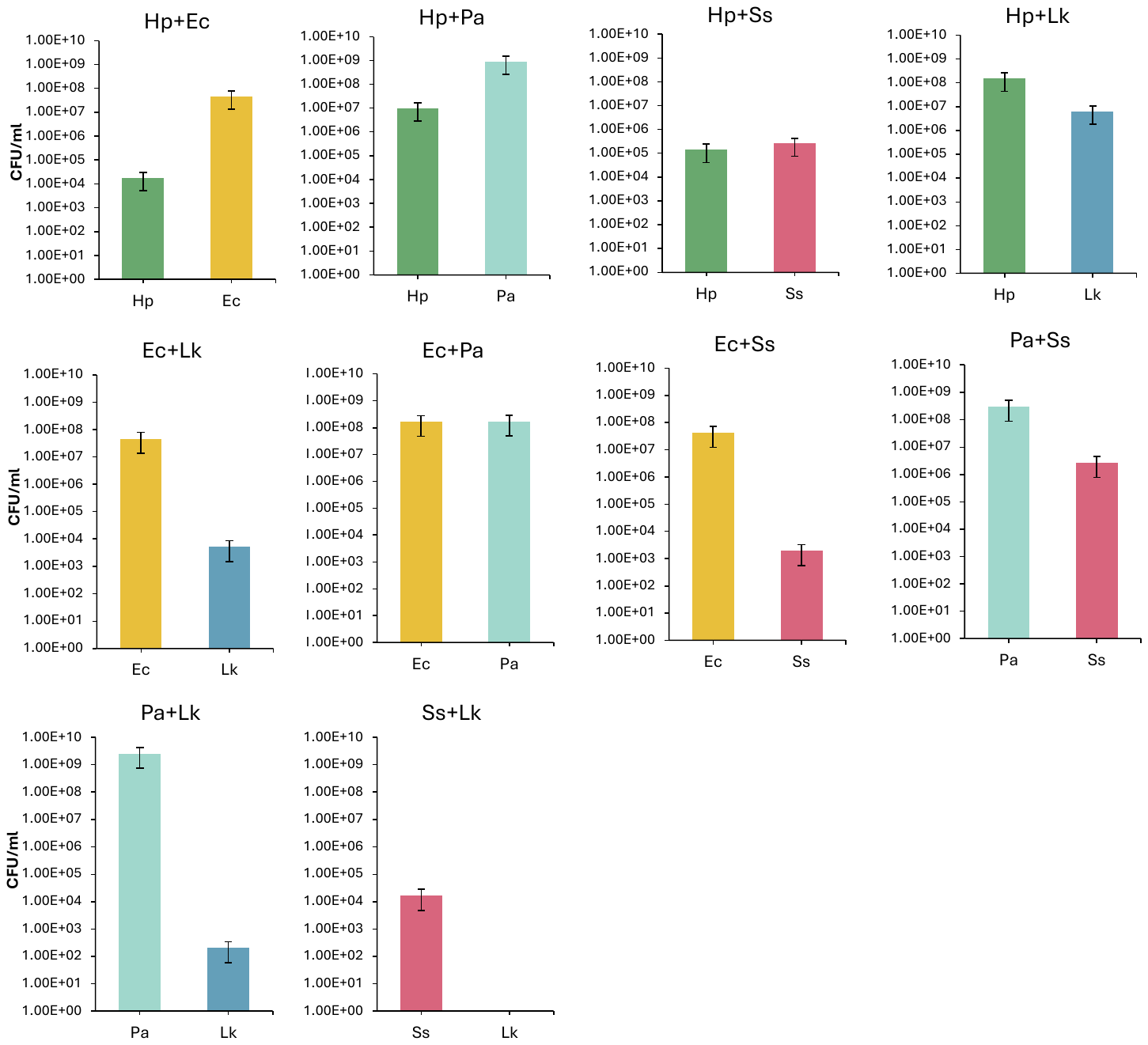

**Supplemental Figure 4.** CFUs of all duo-cultures after 24h co-culture in Ham’s F12.

pH=6.0. Hp=*H. pylori*, Ec=*E. coli*, Pa=*P. aeruginosa*, Ss=*S. salivarius*, Lk=*L. kalixensis*. The bars represent an average of three replicates. Error bars represent +/- SEM.

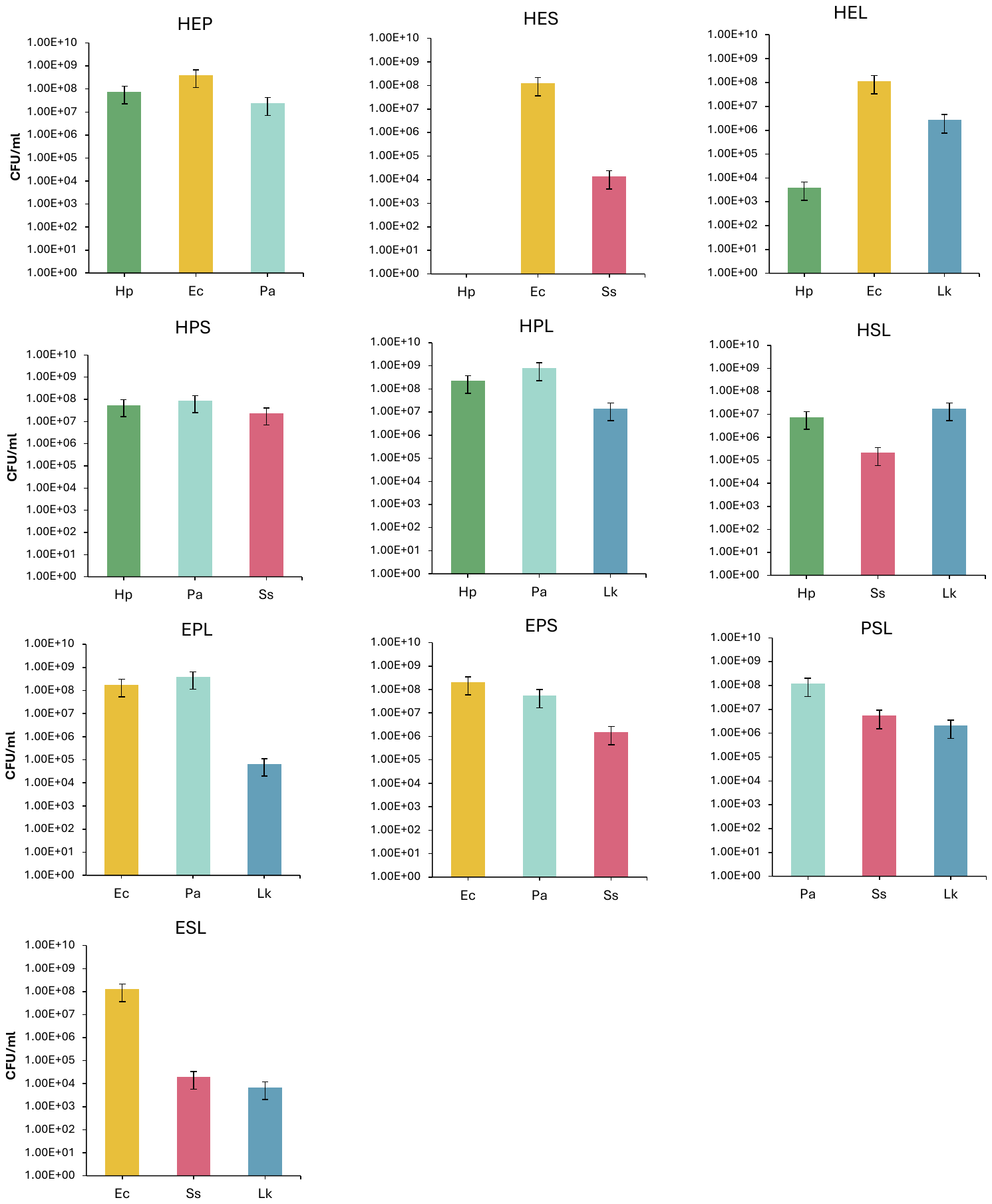

**Supplemental Figure 5:** CFUs of all trio-cultures after 24h co-culture in Ham’s F12.

pH=6.0. Hp=*H. pylori*, Ec=*E. coli*, Pa=*P. aeruginosa*, Ss=*S. salivarius*, Lk=*L. kalixensis*. The bars represent an average of three replicates. Error bars represent +/- SEM.

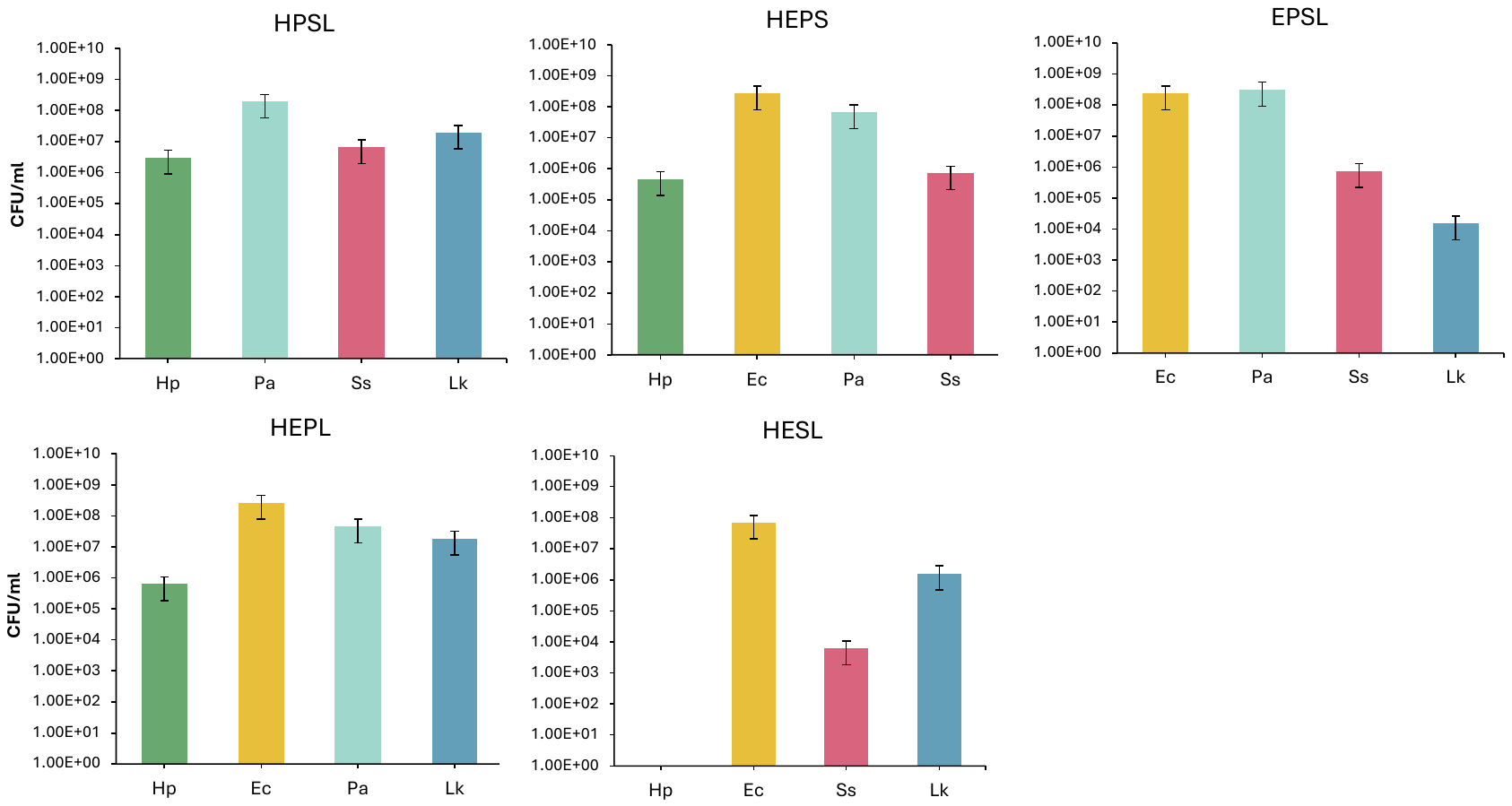

**Supplemental Figure 6:** CFUs of all quartet-cultures after 24h co-culture in Ham’s F12.

pH=6.0. Hp=*H. pylori*, Ec=*E. coli*, Pa=*P. aeruginosa*, Ss=*S. salivarius*, Lk=*L. kalixensis*. The bars represent an average of three replicates. Error bars represent +/- SEM.

|  | | **# of used carbon sources** |  | | | | |
| --- | --- | --- | --- | --- | --- | --- | --- |
|  | **Hp** | 10 | 1.00 | 0.90 | 1.00 | 0.10 | 0.10 |
|  | **Ec** | 12 | 0.75 | 1.00 | 0.92 | 0.08 | 0.08 |
|  | **Pa** | 19 | 0.53 | 0.58 | 1.00 | 0.05 | 0.05 |
|  | **Ss** | 1 | 1.00 | 1.00 | 1.00 | 1.00 | 1.00 |
|  | **Lk** | 1 | 1.00 | 1.00 | 1.00 | 1.00 | 1.00 |
|  |  | | **Hp** | **Ec** | **Pa** | **Ss** | **Lk** |
|  |  |  | **Proportion of Y-axis strain’s carbon sources also used by strains on X-axis** | | | | |

**Supplemental Figure 7**: Carbon source utilization overlap in Ham’s F12. A total of 22 out of the 25 possible carbon sources in Ham’s F12 were included in the phenotype microarray plates. The total number out of the 22 tested carbon sources a strain can utilize is stated in the leftmost column. The numbers represent the proportion of carbon sources the strains on the X-axis can utilize out of all the carbon sources the strain on the Y-axis utilizes. The averages of three independent replicates performed over at least two days were used for calculations.
